# Comparative transcriptome database for *Camellia sinensis* reveals genes important for flavonoid synthesis in tea plants

**DOI:** 10.1101/2024.01.25.577142

**Authors:** Xinghai Zheng, Zahin Mohd Ali, Peng Ken Lim, Marek Mutwil, Yuefei Wang

**Author notes:** Corresponding author: Marek Mutwil, School of Biological Sciences, Nanyang Technological University, 60 Nanyang Drive, Singapore, 637551, Singapore,; Wang Yuefei, Tea Research Institute, Zhejiang University, Hangzhou, Zhejiang, 310058, China.

## Abstract

Tea, as one of the most popular beverages in the world, possesses a plethora of secondary metabolites that are beneficial to human health. Tea plants (*Camellia sinensis*) exhibit rich genetic diversity, where different cultivars can vary significantly in terms of yield, adaptability, morphology, and composition of secondary metabolites. Many tea cultivars have been the subject of much research interest, which have led to the accumulation of publicly available RNA-seq data. As such, it has become possible to systematically summarize the characteristics of different cultivars at the transcriptomic level, identify valuable functional genes, and infer gene functions through co-expression analysis. Here, the transcriptomes of 9 cultivars of *Camellia sinensis* were assembled and combined with the coding sequences of 13 cultivars of *Camellia sinensis* to study the differences and similarities of gene expression and biological functions among cultivars. To give access to this data, we present TeaNekT (https://teanekt.sbs.ntu.edu.sg/), a web resource that provides user-friendly tools and interactive visualizations that facilitates the prediction of gene functions of various tea cultivars. We used TeaNekT to perform cross-cultivar comparison of co-expressed gene neighborhoods, clusters, and tissue-specific gene expression. We show that the members of the chalcone synthase (CHS) gene family, important for flavonoid synthesis, exhibit the highest expression variability, specific expression in leaves and buds, and significant modulation by methyl jasmonate (MeJA) treatment. By using comparative co-expression tools of TeaNekT, we identified multiple conserved genes involved in flavonoid biosynthesis among cultivars that have not been previously studied, warranting further research.

## Introduction

As one of the most popular non-alcoholic beverages worldwide, tea contains a wide range of secondary metabolites beneficial to human health, such as polyphenols, alkaloids, and amino acids (Wang et al., 2022). With the unique qualities of tea plants (*Camellia sinensis*) attracting an increasing number of researchers to study various aspects of tea plants, such as secondary metabolites and phenotypic variations (Zhao et al., 2022; Wang et al., 2022; Liao et al., 2022), the omics dataset of *Camellia sinensis* has also become increasingly extensive. This has led to the use of systems biology approaches on sequencing data hosted on public databases (Tai et al., 2018; Xia et al., 2020; Zhao et al., 2021), such as gene co-expression analysis, becoming a trend in analyzing omics data of *Camellia sinensis*, providing tea researchers with a more macroscopic and comprehensive perspective. Researchers have further downloaded large-scale transcriptome data of tea plants and created a more systematic and comprehensive co-expression database TeaCoN (http://teacon.wchoda.com) (Zhang et al., 2020).

The tea plant (*Camellia sinensis*) possesses a diverse range of germplasm resources (Chen et al., 2007). Different cultivars of *Camellia sinensis* are each prized for certain desirable qualities in their own right and exhibit significant differences in plant morphology, leaf characteristics, growth habits, adaptability, and secondary metabolites (Chen et al., 2012; Zhao et al., 2022). For instance, temperature-sensitive albino cultivar “Anji Baicha” and light-sensitive yellowing cultivar “Huangjinya” have attracted much attention due to their unique leaf color and secondary metabolite content (Zhang et al., 2020). The new shoots of “Anji Baicha” cultivar are highly sensitive to cold temperatures and undergo a color transformation as the temperature gradually warms up in early spring (Cheng et al., 1999; Li, 2002). Meanwhile, the amino acid content in dry tea made from the shoots of “Anji Baicha” during the white stage can reach up to 6%, three times higher than that of ordinary cultivars (2%) (Li et al., 1996; Xiong et al., 2013). Additionally, for example, “Zijuan” is a unique cultivar of purple leaf tea, characterized by its purple stems, leaves, and buds. It is a mutant of *Camellia sinensis* var. *assamica* and contains abundant anthocyanins in its leaves (Yang et al., 2009).

Given that the significant differences among different cultivars may arise from variations in coding sequences or expression patterns, it is necessary to start by performing transcriptome assembly of these cultivars to better understand the genetic basis for these metabolic and physiological differences. Although researchers have already studied the differences between different *Camellia sinensis* cultivars through transcriptome assembly and investigated the expansion and evolution of important gene families in tea plants (Kong et al., 2022; Wu et al., 2022), there has been no research that combines assembled transcriptomes with co-expression analysis to further explore the biological functions of genes involved in specific key mechanisms of tea plants.

It has been well-established that functionally related genes often exhibit similar gene expression patterns (Zhou et al., 2002). Therefore, by identifying gene clusters with highly similar expression patterns, we can identify functionally related genes and infer the functions of unknown genes based on the functions of their neighboring genes (Usadel et al., 2009). Moreover, the conservation of functionally related genes among different cultivars would further confirm their importance (Hansen et al., 2014). In the view of this, we are set to construct a co-expression analysis tool that can effectively mine predictive information on gene function and regulation from transcriptomic data. To this end, we utilize CoNekT (Co-expression Network Toolkit) (Proost and Mutwil, 2018), a popular framework that has been used to construct comparative transcriptomic databases for plants and species from other kingdoms (Ng et al., 2020; Tan and Mutwil, 2020; Lim et al., 2020; Lim et al., 2022; Villanueva et al., 2022).

In this study, the transcriptomes of 9 *Camellia sinensis* cultivars were individually assembled. Then, the coding sequences of 13 *Camellia sinensis* cultivars were annotated and subjected to phylogenetic analysis. The expression profiles of these 13 *Camellia sinensis* cultivars were calculated and used for constructing co-expression networks. All transcriptome data, including annotations and co-expression networks, of the 13 *Camellia sinensis* cultivars were imported into CoNekT, resulting in the creation of TeaNekT (Co-expression Network Toolkit for tea plants) (https://teanekt.sbs.ntu.edu.sg/). The phylogenetic analysis of the 13 cultivars successfully reconstructed their relationships. In the study on gene expression levels related to flavonoid synthesis, we observed significant variation in the expression levels of genes involved in the phenylpropanoid pathway and flavonoid pathway in response to experimental treatments across several cultivars. With the assistance of TeaNekT, a co-expression cluster named “Cluster_126” containing multiple genes from the flavonoid pathway was discovered in the “Jinxuan” cultivar. The expression profile of this cluster exhibited a high sensitivity to methyl jasmonate (MeJA) treatment, and many genes within the cluster showed conserved co-expression relationships across different cultivars. Additionally, several genes from the flavonoid pathway within this cluster, such as the CHS gene family, exhibited specific expression patterns in the leaves and buds.

## Methods

### Data source and sample metadata annotation

By searching and filtering using the keyword “Camellia sinensis” in the NCBI SRA database, a total of 861 RNA-Seq raw reads from 13 *Camellia sinensis* cultivars were obtained. Initial annotations of these RNA-Seq raw data were performed using the metadata fields in the NCBI SRA database, including cultivar, plant tissue, sampling age, and experimental treatments. Subsequently, the corresponding original papers for each RNA-Seq data were searched and retrieved to further supplement and correct the annotation information (Table S1).

For the convenience of calculating the specificity measure (SPM) in the TeaNekT database for all plant tissues, only six terms were retained: “leaf, bud”, “root”, “stem”, “flower”, “fruit” and “seed” (e.g., descriptions like “two leaves and a bud” were classified under “leaf, bud”). For the control group, samples in each experiment or samples directly collected without any treatment, they were uniformly labeled as “no treatment” in the experimental treatment column. For the samples with missing annotations in the metadata fields of the NCBI SRA database and could not be found in the retrieved original papers were labeled as “missing”.

### Genome-guided transcriptome assembly and completeness assessment

The coding sequences of the cultivar “Shuchazao” was downloaded from TPIA (http://tpia.teaplant.org) (Xia et al., 2019), and the coding sequences of the cultivars “Huangdan”, “Longjing 43” and “Yunkang 10” were downloaded from TeaPGDB V1.0 (http://eplant.njau.edu.cn/tea) (Lei et al., 2021). The coding sequences of the other 9 cultivars of *Camellia sinensis* were obtained using the following genome-guided transcriptome assembly method.

Firstly, several RNA-Seq raw data containing different metadata entries were selected from each cultivar for transcriptome assembly. The RNA-Seq raw data were processed using the fastp tool (Chen et al., 2018) to remove adapters and low-quality reads, resulting in high-quality clean data. Then, the HISAT2 sequence alignment software (Kim et al., 2015) was used to align the clean data reads to the reference genome of “Shuchazao” downloaded from TPIA (http://tpia.teaplant.org) (Xia et al., 2019), generating BAM files. The BAM files were then assembled using the TRINITY tool with default parameters (Haas et al., 2013), resulting in the assembled transcripts. Next, the TransDecoder tool with default parameters (https://transdecoder.github.io/) was used to identify and predict open reading frames (ORFs) in the transcripts assembled by TRINITY, identifying potential protein-coding sequences. Finally, the CD-HIT tool with default parameters (Fu et al., 2012) was used to reduce unavoidable redundancy introduced during the assembly process, resulting in the final CDS file, i.e., the coding sequences (Table S2).

The completeness of the transcriptome assembly was evaluated using the BUSCO tool (Simão et al., 2015) based on the eudicots_odb10 dataset, which represents the dataset of eudicot plants, for both the assembled transcripts generated by the TRINITY tool for the 9 *Camellia sinensis* cultivars and the coding sequences of the 13 *Camellia sinensis* cultivars.

### Conserved genes and cultivar-specific genes identification

Firstly, the coding sequences of 13 cultivars of *Camellia sinensis* were aligned using the OrthoFinder tool with default parameters (Emms & Kelly, 2019), and the orthologous groups (OG) among these cultivars were constructed. Next, coding sequences present in the orthologous groups containing coding sequences from all 13 cultivars were extracted as conserved genes, while coding sequences present only in the orthologous groups of a single cultivar were extracted as cultivar-specific genes.

### Gene annotation and functional enrichment analysis

The coding sequences of the 13 *Camellia sinensis* cultivars were annotated for biological functions using the Mercator v4 2.0 online tool (Lohse et al., 2014) (Table S3). The InterProScan tool (Quevillon et al., 2005) was used to obtain protein Pfam domains and Gene Ontology (GO) terms for each gene from the coding sequences of the 13 *Camellia sinensis* cultivars (Table S3). The KEGG annotation of the coding sequences of the 13 *Camellia sinensis* cultivars was performed using the BlastKOALA online tool (Kanehisa et al., 2016) available at https://www.kegg.jp/blastkoala/ (Table S3). The iTAK tool with default parameters (Zheng et al., 2016) was used to predict and classify transcriptional factors from the coding sequences of the 13 *Camellia sinensis* cultivars (Table S3).

To perform functional annotation on the conserved genes and cultivar-specific genes, the hypergeometric test was conducted using the hypergeom tool from the scipy.stats package (Hahne et al., 2008). This test compared the genes in each conserved gene and cultivar-specific genes with the genes associated with each annotation term. For the mapman annotation, only annotation terms with more than 100 genes and the most detailed classification entries were selected. Then, the p-values of all annotation terms corresponding to each conserved genes and cultivar-specific genes were corrected using the fdrcorrection tool from the statsmodels.stats.multitest package (Benjamini and Hochberg, 1995; Nazer et al., 2018) to obtain the False Discovery Rate (FDR) values (Table S4). Annotation terms with FDR values less than or equal to 0.05 were considered significant for functional annotation of the conserved genes and cultivar-specific genes.

### Phylogenetic analysis of *Camellia sinensis* cultivars

A set of single-copy orthologous genes containing a single orthologous gene from each of the 13 *Camellia sinensis* cultivars was extracted from the conserved genes for phylogenetic analysis.

To construct the phylogenetic relationships among *Camellia sinensis* cultivars, the amino acid sequences of each single-copy orthologous gene pair were aligned using the MUSCLE tool with default parameters (Edgar, 2004). The TRIMAL tool (Capella-Gutierrez et al., 2009) was then used to remove poorly aligned regions. Based on the trimmed alignment files of each single-copy orthologous gene pair, phylogenetic trees were constructed using the RAXML tool (Stamatakis, 2014), with “Yunkang 10” serving as the outgroup. Subsequently, all the phylogenetic tree files of the single-copy orthologous gene pairs were merged into a supermatrix file, and the ASTRAL tool (Zhang et al., 2018) was used to construct the phylogenetic tree of the 13 *Camellia sinensis* cultivars.

### Gene expression levels and expression variation coefficients calculation

The clean data reads were pseudo-aligned to the coding sequences of the 13 *Camellia sinensis* cultivars using the kallisto tool (Bray et al., 2016). This allowed for the quantification of the transcripts per million (TPM) values for all genes in each sample, serving as a measure of gene expression levels. Additionally, the percentage of sequencing reads aligned to the coding sequences was calculated as the pseudo-alignment read percentage, providing an assessment of the alignment efficiency of the transcriptomic sequencing data (Table S1) (Figure S1). Subsequently, the coefficient of variation (CV) for the expression levels of each gene across all samples was calculated as an indicator of gene expression variability within each *Camellia sinensis* cultivar (Table S1).

### Co-expression networks and TeaNekT database construction

First, an expression profile was constructed for the 13 *Camellia sinensis* cultivars. Then, a co-expression network was built for the *Camellia sinensis* cultivars using the Highest Reciprocal Rank metric (Mutwil et al., 2010). The CoNekT database framework, with default settings, was used to populate the database with information such as gene expression levels, gene annotations, orthologous gene sets, and phylogenetic trees for the 13 *Camellia sinensis* cultivars (Proost and Mutwil, 2018). This resulted in the construction of TeaNekT (https://teanekt.sbs.ntu.edu.sg/), an online *Camellia sinensis* database equipped with co-expression clusters that can identify co-expressed genes, gene families, and enrich specific biological functions. In TeaNekT database, the Heuristic Cluster Chiseling Algorithm (HCCA) (Mutwil et al., 2010) was employed to generate co-expression clusters for the 13 *Camellia sinensis* cultivars, with a limit of 100 genes per cluster.

## Results

### Information on selected *Camellia sinensis* cultivars

This study selected a total of 13 *Camellia sinensis* cultivars, including “Anji Baicha”, “Echa 1”, “Fuding Dabaicha”, “Huangdan”, “Huangjinya”, “Jinxuan”, “Longjing 43”, “Longjing Changye”, “Shuchazao”, “Tieguanyin”, “Zhongcha 108”, “Yunkang 10” and “Zijuan” for transcriptome assembly and the TeaNekT database construction. These cultivars were chosen since they have more RNA-Seq data available in NCBI (at least 30 RNA-Seq data) compared to other *Camellia sinensis* cultivars. This not only indicates their greater research value but also ensures sufficient data support for this study and the database.

Among these 13 *Camellia sinensis* cultivars, 11 of them belong to *Camellia sinensis* var. *sinensis*, while two cultivars, “Yunkang 10” and “Zijuan”, belong to *Camellia sinensis* var. *assamica* (Table 1). Only the coding sequences of “Shuchazao” was found in the TPIA database, and the coding sequences of “Yunkang 10”, “Longjing 43”, and “Huangdan” were found in TeaPGDB V1.0 database, while the coding sequences of the other 9 *Camellia sinensis* cultivars were assembled by this study.

**Table 1.**
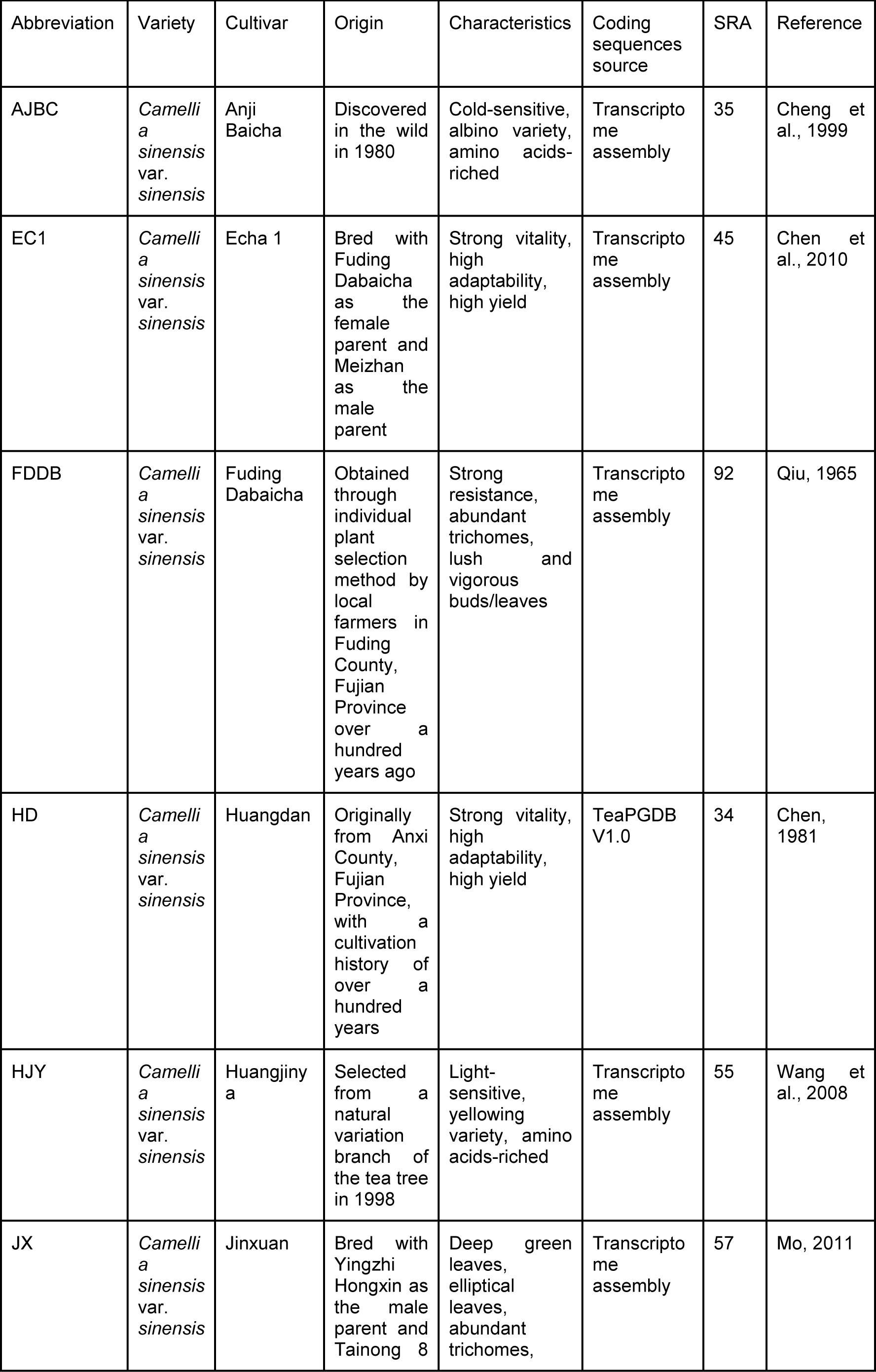
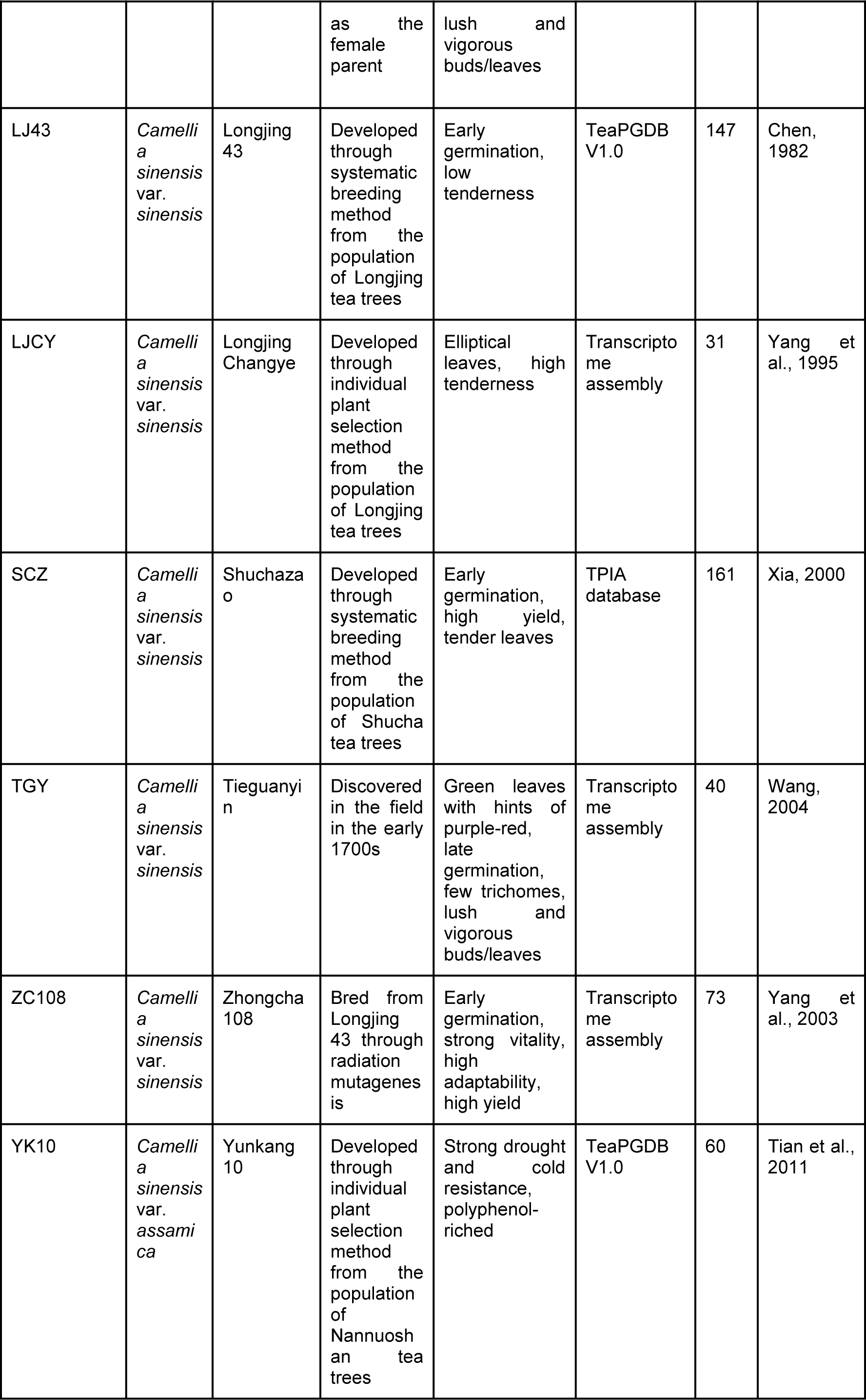
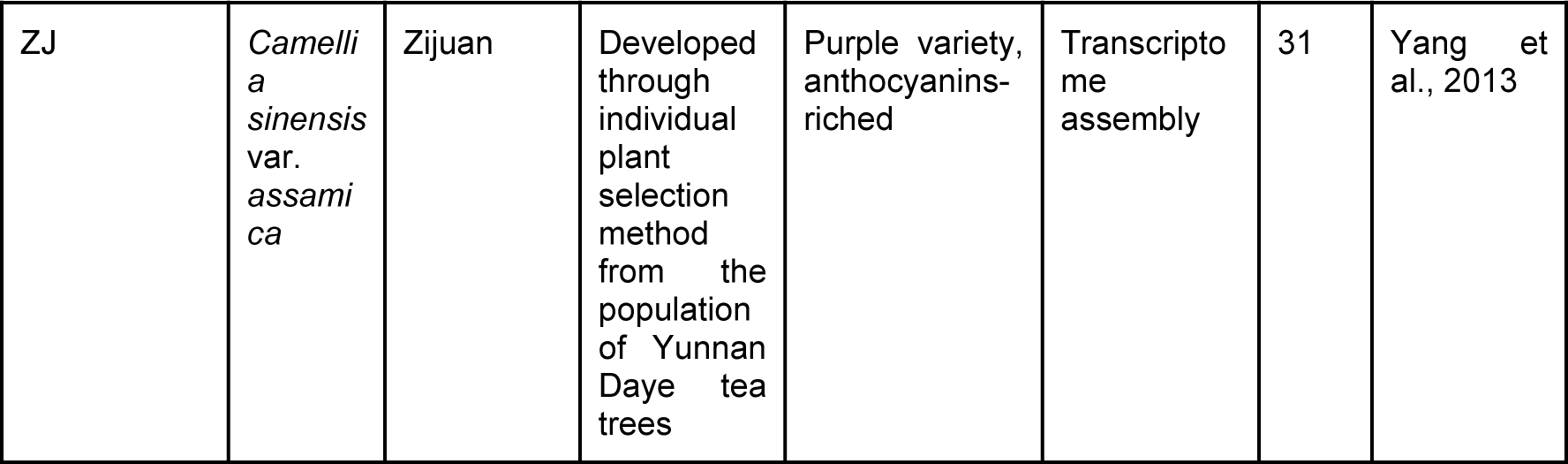
Overview of 13 *Camellia sinensis* Cultivars.

### Genome-guided transcriptome assembly yielded high completeness of coding sequences

Through downloading and genome-guided transcriptome assembly, coding sequences from 13 *Camellia sinensis* cultivars were obtained (Table S2). The transcriptome assembly of the 9 *Camellia sinensis* cultivars was carried out based on the following pipeline (Figure 1A). Subsequently, the coding sequences of the 13 *Camellia sinensis* cultivars were annotated, subjected to phylogenetic analysis, expression quantification, co-expression network analysis, and construction of the TeaNekT database.

**Figure 1.**
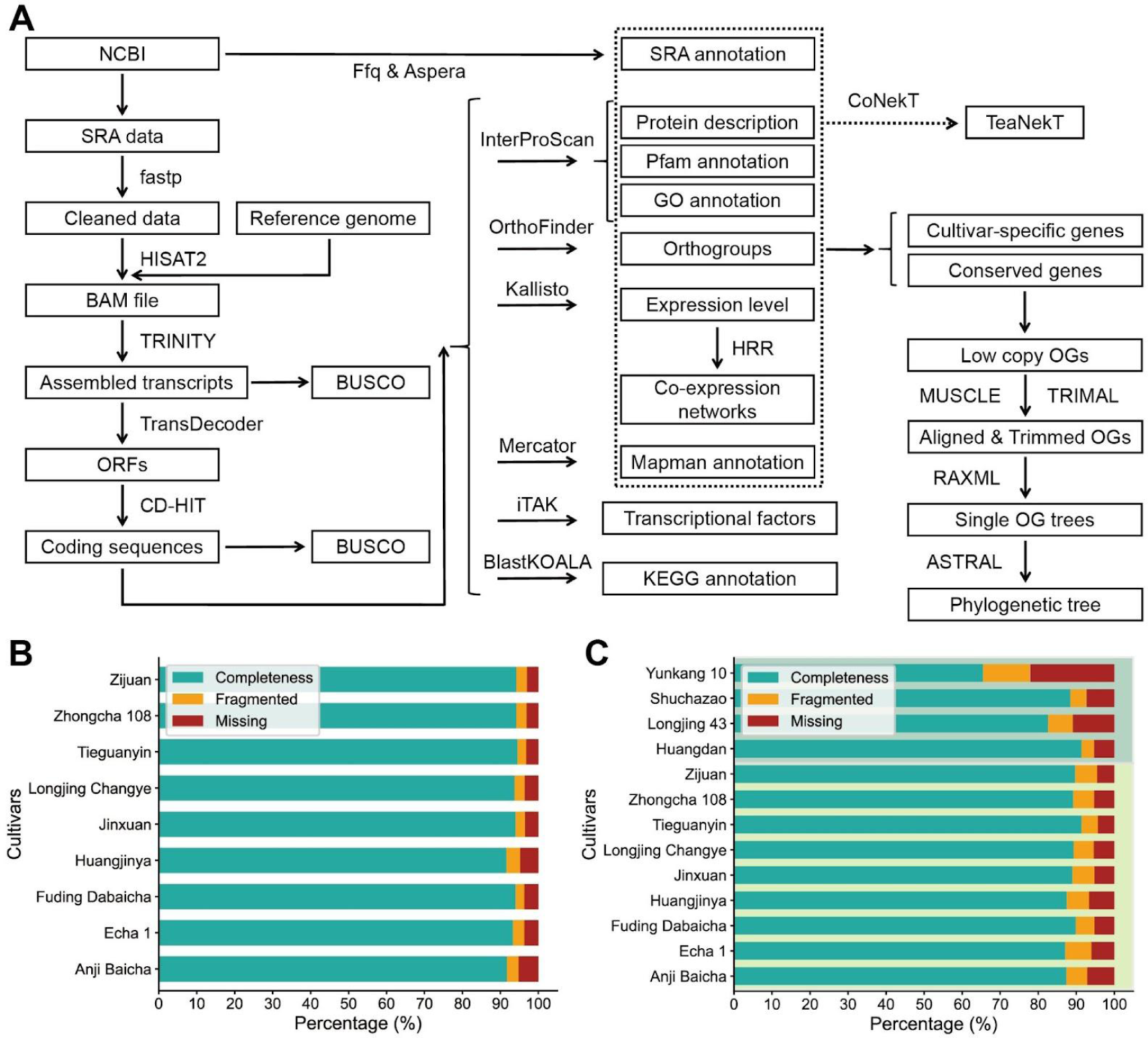
Pipeline of the genome-guided transcriptome assembly and TeaNekT database construction, along with the completeness assessment of the assembled transcripts and coding sequences. (A) Pipeline of the genome-guided transcriptome assembly and TeaNekT database construction. (B) Completeness assessment of the assembled transcripts by BUSCO. (C) Completeness assessment of the coding sequences by BUSCO. Cultivars with coding sequence downloaded from database and cultivars with coding sequence self-assembled are distinguished by different colored background panels (Top: Downloaded from database, Bottom: Self-assembled).

During the assembly process, the assembled transcripts generated by TRINITY yielded approximately 360,000 to 670,000 sequences, many of which could represent non-coding sequences or redundant transcripts (Table 2). To obtain coding transcripts, TransDecoder was used to filter and obtain approximately 110,000 to 180,000 coding sequences. Finally, CD-HIT was employed to remove duplicate sequences, resulting in approximately 49,000 to 67,000 coding sequences. The quality of the assembly for both the assembled transcripts of the 9 cultivars and the coding sequences of the 13 cultivars were assessed using BUSCO.

**Table 2.** Change in the number of sequences from assembled transcripts to the coding sequences.

| Cultivars | AJBC | EC1 | FDDDB | HJY | JX | LJCY | TGY | ZC108 | ZJ |
| --- | --- | --- | --- | --- | --- | --- | --- | --- | --- |
| Assembled transcripts | 662791 | 587518 | 533846 | 508150 | 597634 | 591662 | 361842 | 522222 | 420751 |
| ORFs | 172926 | 176696 | 151144 | 146106 | 179914 | 174391 | 111671 | 160814 | 132113 |
| Coding sequences | 64925 | 66747 | 60805 | 58213 | 66303 | 65999 | 49257 | 61055 | 59405 |

In the BUSCO results, we observed that the assembled transcripts assemblies of the 9 *Camellia sinensis* cultivars exhibit completeness values of over 90% (Figure 1B). Furthermore, the completeness values of the coding sequences are also above 85% (Figure 1C). These values are comparable to the quality of “Shuchazao” and “Huangdan”, and significantly higher than that of “Longjing 43” and “Yunkang 10”. This indicates that the transcriptome assembly in this study is reliable.

### Comparison of CDS obtained from the genome and transcriptome assembly

To present the basic characteristics of the coding sequences of 13 *Camellia sinensis* cultivars, we examined the number of transcripts, GC content, transcript length distribution, conserved genes and cultivar-specific genes ratio, and annotation rate.

The number of transcripts of *Camellia sinensis* cultivars ranges from approximately 30,000 to 70,000 (Figure 1A). In this study, the assembled coding sequences generally contained a higher number of coding sequences compared to the coding sequences available in the TPIA and TeaPGDB V1.0 database. Specifically, the “Yunkang 10” and “Longjing 43” cultivars downloaded from the database had the fewest number of coding sequences, with 36,951 and 33,556 coding sequences, respectively. The GC content of all *Camellia sinensis* cultivar coding sequences remains relatively stable, fluctuating around 44% (Figure 2A).

**Figure 2.**
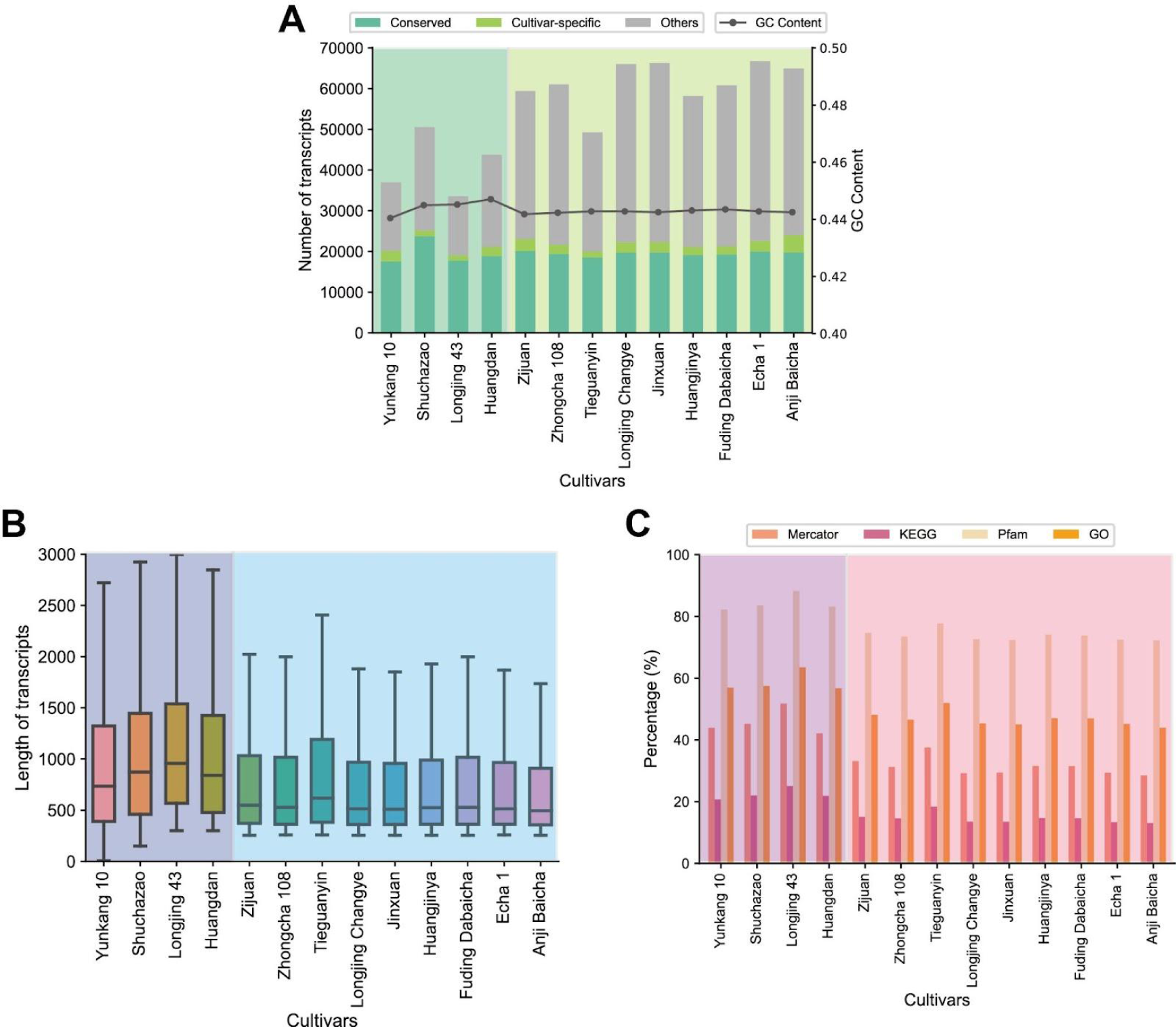
Characteristics of 13 *Camellia sinensis* cultivars’ coding sequence. Cultivars with coding sequences downloaded from database and cultivars with coding sequences self-assembled are distinguished by different colored background panels (Left: Downloaded from database, Right: Self-assembled). (A) GC content and proportion of conserved genes and cultivar-specific genes. (B) Length distribution. (C) Proportion of annotated genes.

In this study, conserved genes are those that exist in the coding sequences of homologous groups containing 13 *Camellia sinensis* cultivars, while cultivar-specific genes refer to those that exist in the coding sequences of homologous groups of only one *Camellia sinensis* cultivar. In all *Camellia sinensis* cultivar coding sequences, the number of conserved genes remains stable, fluctuating around 20,000 (Figure 2A). The number of cultivar-specific genes does not exceed 5,000 in any cultivar. The number of conserved genes remains consistent across the cultivars, while the number of non-conserved genes (the collection of “Others” and “Cultivar-specific” genes) shows significant variation among the cultivars.

In all *Camellia sinensis* cultivar coding sequences, the median transcript length ranges from 500 to 1000 (Figure 2B) (Figure S2). The median transcript length values of the assembled coding sequences are generally lower than those of the coding sequences available in the TPIA and TeaPGDB V1.0 database.

Annotations were performed on all *Camellia sinensis* cultivar coding sequences, including KEGG annotation, GO annotation, Pfam annotation, and Mercator annotation (Figure 2C). We found that approximately 70% to 85% of the coding sequences had Pfam annotations, approximately 40% to 60% had GO annotations, approximately 30% to 50% had Mercator annotations, and approximately 10% to 20% had KEGG annotations. The annotation rates of the assembled coding sequences were generally lower than those of the coding sequences available in the TPIA and TeaPGDB V1.0 database.

### Phylogenetic analysis revealed the interrelationships among *Camellia sinensis* cultivars

To analyze the phylogenetic relationship and transcriptomic similarity among *Camellia sinensis* cultivars, a phylogenetic analysis was conducted on coding sequences of 13 *Camellia sinensis* cultivars (Figure 3). ASTRAL utilizes maximum likelihood estimation and the bootstrap method to calculate branch support values. In this phylogenetic tree, branch support values are typically close to 1. A high branch support value (close to 1) indicates that the branch is well-supported across multiple datasets, allowing for confident inference of its presence and position. We found that the phylogenetic relationships and order of the 13 cultivars were consistent with the origin and characteristics of the cultivars listed in Table 1. This further confirms the high reliability of the assembled coding sequences.

**Figure 3.**
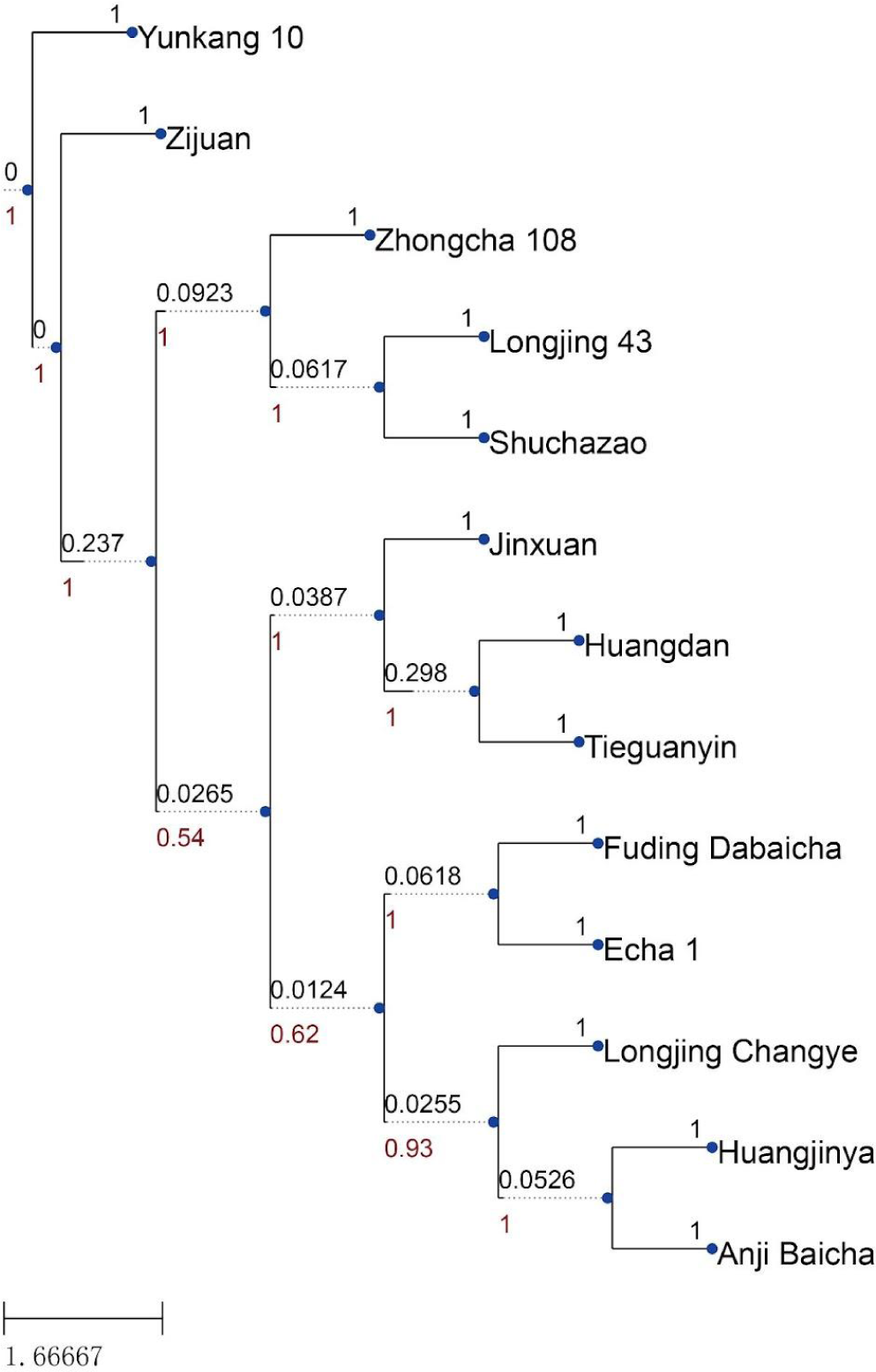
Phylogenetic tree of 13 *Camellia sinensis* cultivars. Bootstrap values and posterior probabilities are colored and displayed on the branches of the phylogeny (Red for branch support values and black for branch length values).

The phylogenetic order of the 13 *Camellia sinensis* cultivars, displayed from top to bottom, is as follows: “Yunkang 10” and “Zijuan”, as more primitive var. *assamica* cultivars, diverge from the other cultivars in the earliest branch. “Zhongcha 108”, as an artificially irradiated variant of “Longjing 43”, is grouped in a neighboring branch with “Longjing 43”. “Echa1”, a hybrid cultivar derived from the cross between “Fuding Dabaicha” as the maternal parent and “Meizhan” as the paternal parent, is placed in the same branch as “Fuding Dabaicha”. Finally, “Anji Baicha” and “Huangjinya”, two unique colored tea cultivars that undergo leaf color changes, are sensitive to the environment and rich in amino acids, are grouped together in a separate branch.

### Study on the synthesis pathways of flavonoids in *Camellia sinensis* identified several cultivars with significant expression level variations

Tea leaves contain a large amount of flavonoids, which are the most important class of secondary metabolites in tea and one of the key factors determining its unique quality (Wang et al., 2022). Therefore, studying the key genes involved in flavonoid synthesis pathways can provide us with more insights into the formation of the unique qualities of tea. The synthesis pathway of flavonoids can be divided into three steps: shikimic acid pathway, phenylpropanoid pathway, and flavonoid pathway (Figure 4A). Among them, the latter two steps are particularly crucial for the synthesis of flavonoids.

**Figure 4.**
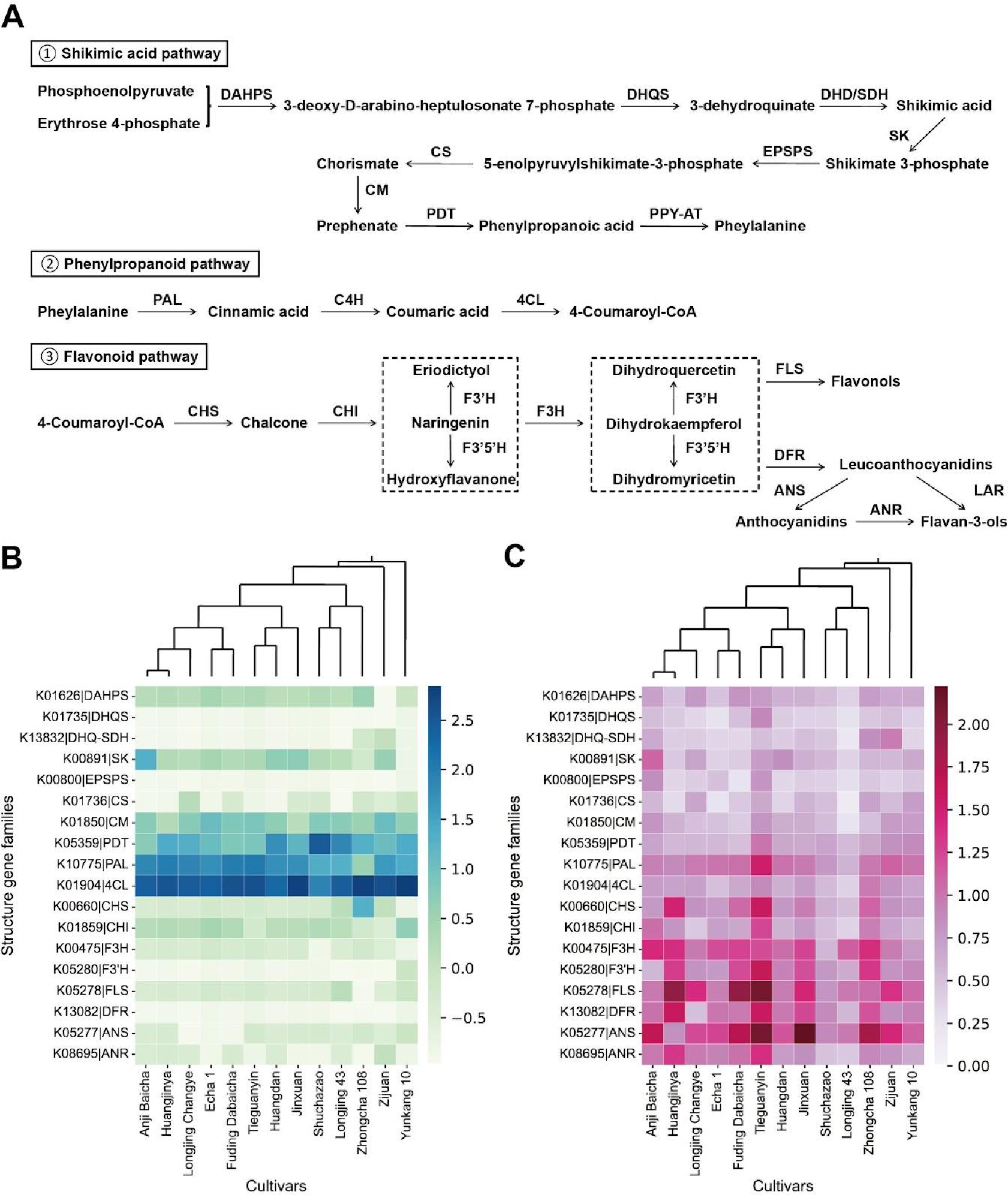
Copy number and median coefficient of variation (CV) of expression levels among structural gene families involved in the synthesis pathways of flavonoid in *Camellia sinensis*. DAHPS: 3-deoxy-D-arabinoheptulosonate-7-phosphate synthase; DHQS: 3-dehydroquinate synthase; DHD: 3-dehydroquinate synthase; SDH: dehydrogenase; SK: shikimate kinase; EPSPS: 5-enolpyruvylshikimate-3-phosphate synthase; CS: chorismate synthase; CM: chorismate mutase; PDT: prephenate dehydratase; PPY-AT: phenylpyruvate transaminase; PAL: phenylalanine ammonialyase; C4H: cinnamate 4-hydroxylase; 4CL: p-coumaroyl-CoA ligase; CHS: chalcone synthase; CHI: chalcone isomerase; F3H: naringenin 3-dioxygenase; F3’H: flavonoid 3’-hydroxylase; F3’5’H: flavonoid 3’,5’-hydroxylase; FLS: flavonol synthase; DFR: dihydroflavonol 4-reductase; ANS: anthocyanidin synthase; ANR: anthocyanidin reductase; LAR: leucoanthocyanidin reductase. (A) The three-stage flavonoid biosynthesis pathway in *Camellia sinensis*. (B) Copy number. The color of the cells represents the copy number. The sorting of biological functional terms is based on the hierarchical clustering of the copy number, while the sorting of cultivars is based on the phylogenetic tree. (C) Median coefficient of variation (CV) of expression levels. The color of the cells represents the median coefficient of variation (CV) of expression levels. The sorting of transcription factor families is based on the hierarchical clustering of the median coefficient of variation (CV) of expression levels, while the sorting of cultivars is based on the phylogenetic tree.

In the study of structural gene family size, we found that almost all cultivars have a high number of copies for phenylalanine ammonialyase (PAL) and p-coumaroyl-CoA ligase (4CL) structural gene families, which happen to correspond to the genes encoding the major enzymes in phenylpropanoid pathway (Figure 4B).

Based on the fact that numerous researchers have conducted various experimental treatments on different cultivars of tea plants, here we provide a convenient approach to identify which experimental treatments are most sensitive for the structural gene families involved in the synthesis pathways of flavonoids. We plotted the median variation in expression levels of various structural gene families involved in the synthesis pathways of flavonoids across different cultivars under different experimental treatments. We observed that there is a gradual increase in the median expression level variation of gene families from the shikimic acid pathway to the flavonoid pathway (Figure 4C). Particularly, a larger median variation is observed in the phenylpropanoid pathway and flavonoid pathway, where the flavonol synthase (FLS) family and anthocyanidin synthase (ANS) family exhibit extremely high coefficient of variation (CV) in several cultivars. It is conceivable that in tea plant research, researchers tend to apply experimental treatments that result in significant expression differences in pathway genes during the flavonoid pathway.

### TeaNekT assisted in the discovery of a highly sensitive co-expression cluster to methyl jasmonate (MeJA) in “Jinxuan”

The study on the synthesis pathways of flavonoids revealed significant differential expression of a series of genes in the flavonoid pathway caused by experimental treatments in several cultivars, such as “Tieguanyin”, “Jinxuan”, “Huangjinya”, and “Fuding Dabaicha” (Figure 4C). To identify the experimental treatments that cause significant differential expression of genes in the flavonoid pathway, we utilized TeaNekT (https://teanekt.sbs.ntu.edu.sg/) to study the expression profiles of genes in the flavonoid pathway across various cultivars. We discovered that in the cultivar “Jinxuan”, most of the genes related to the flavonoid pathway are co-expressed in the HCCA cluster “Cluster_126” (https://teanekt.sbs.ntu.edu.sg/cluster/view/7475). Therefore, we generated average expression profiles for the co-expression cluster “Cluster_126” and the expression profiles of genes related to the flavonoid pathway in “Cluster_126” (https://teanekt.sbs.ntu.edu.sg/heatmap/cluster/7475) (Figure 5AB). We observed that genes in “Cluster_126” exhibit similar expression patterns and are highly responsive to methyl jasmonate (MeJA) treatment (showing significant upregulation in response to methyl jasmonate (MeJA) treatment). Moreover, their expression is significantly upregulated in regenerating tissues.

The co-expression gene cluster “Cluster_126” consists of 144 genes (https://teanekt.sbs.ntu.edu.sg/cluster/view/7475). The genes in “Cluster_126” are significantly enriched in biological functions like oxidoreductase activity, UDP-glycosyltransferase activity, aromatic amino acid family biosynthetic process, organic cyclic compound biosynthetic process, and aromatic amino acid metabolic process, among others. With the assistance of TeaNekT, we were able to construct a co-expression network for “Cluster_126”, which provided us with a more detailed understanding of the general functions of each gene within “Cluster_126” (https://teanekt.sbs.ntu.edu.sg/cluster/graph/7475) (Figure 5C). The genes in the cluster are related to “External stimuli response” (*PKS4*), “Phytohormone action” (*GASA3*), and seven genes related to “Transcriptional regulation” (*TT8*, *JAI3*, *MYB4*, *GIF3*, *MYB3*, *ATY53* and *JINXUAN17122*). In addition, genes associated with flavonoid synthesis include three phenylalanine ammonialyase (PAL) genes (*JINXUAN26315*, *JINXUAN15354* and *JINXUAN27063*), one cinnamate 4-hydroxylase (C4H) (*JINXUAN33459*), two p-coumaroyl-CoA ligases (4CL) (*JINXUAN42050* and *JINXUAN17222*), two chalcone synthases (CHS) (*JINXUAN00181* and *JINXUAN14849*), one naringenin 3-dioxygenase (F3H) (*JINXUAN60265*), and one dihydroflavonol 4-reductase (DFR) (*JINXUAN43808*).

**Figure 5.**
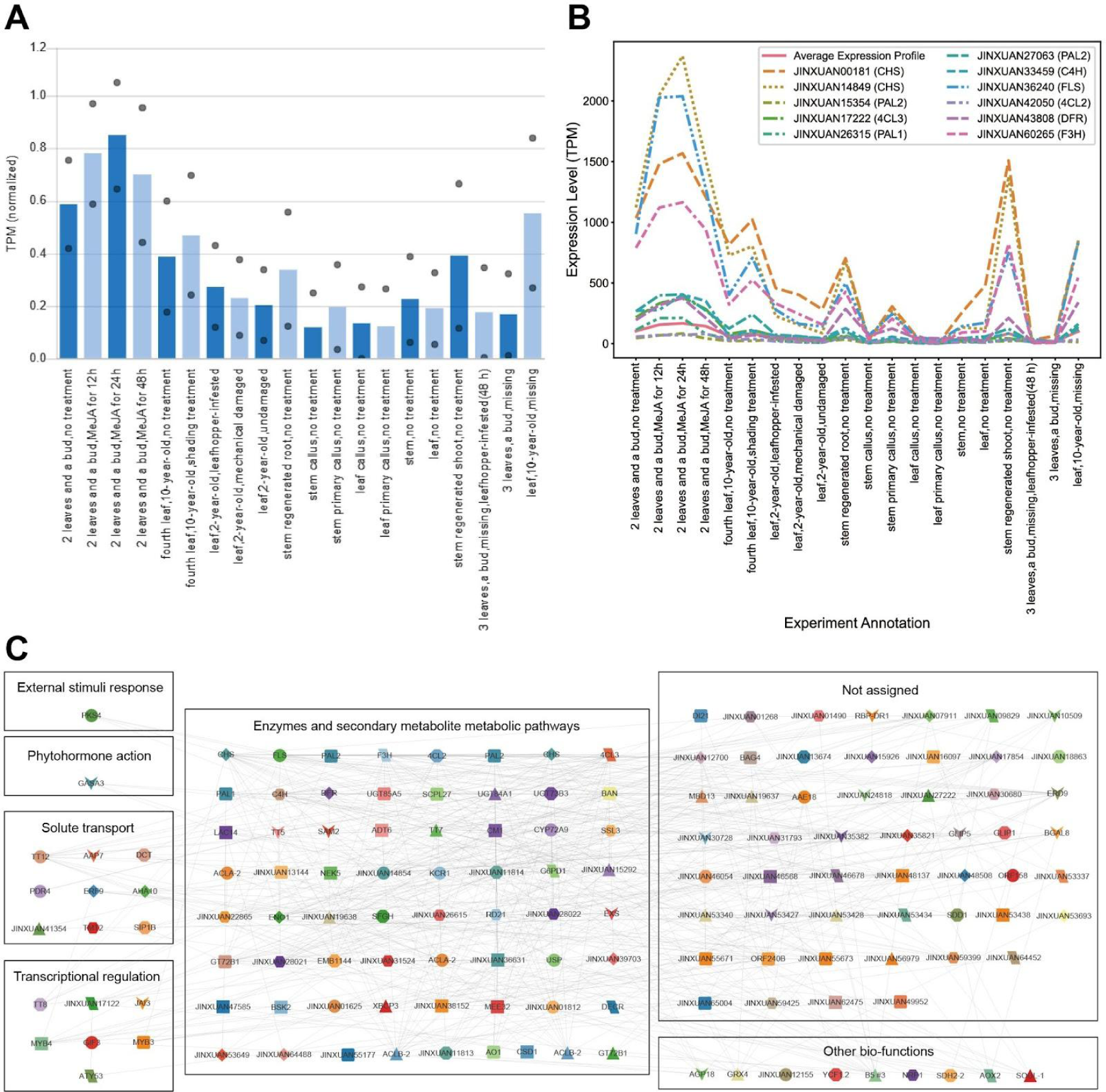
Expression profiles of and the co-expression network of genes in the co-expression cluster “Cluster_126” of the “Jinxuan” cultivar. (A) Average expression profile of the co-expression cluster “Cluster_126” in the “Jinxuan” cultivar from TeaNekT. (B) Average expression profile of the co-expression cluster “Cluster_126” in the “Jinxuan” cultivar and the expression profile of genes related to the flavonoid synthesis pathway. (C) Co-expression gene network of the co-expression cluster “Cluster_126” in the “Jinxuan” cultivar. Nodes, colored shapes, and gray edges represent genes, orthogroups, and co-expressed genes, respectively. Genes of different biological functions are classified into different boxes.

### TeaNekT revealed the tissue specificity and conservation of co-expression relationships of the CHS family

In the expression profile of the co-expression cluster “Cluster_126”, we observed that two genes from the chalcone synthase (CHS) gene family, *JINXUAN00181* and *JINXUAN14849*, exhibit the highest expression variability compared to other genes involved in the flavonoid pathway (except for one FLS gene) (Figure 5B). To further investigate the tissue specificity conservation of the chalcone synthase (CHS) gene family, the phylogenetic tree of the OG0001908 orthogroup, which includes *JINXUAN00181* and *JINXUAN14849* was derived from TeaNekT (https://teanekt.sbs.ntu.edu.sg/tree/view/1908) (Figure 6A). We observed that all the homologous genes within the OG0001908 orthogroup are specifically expressed in leaf and bud, while showing minimal expression in other plant tissues.

**Figure 6.**
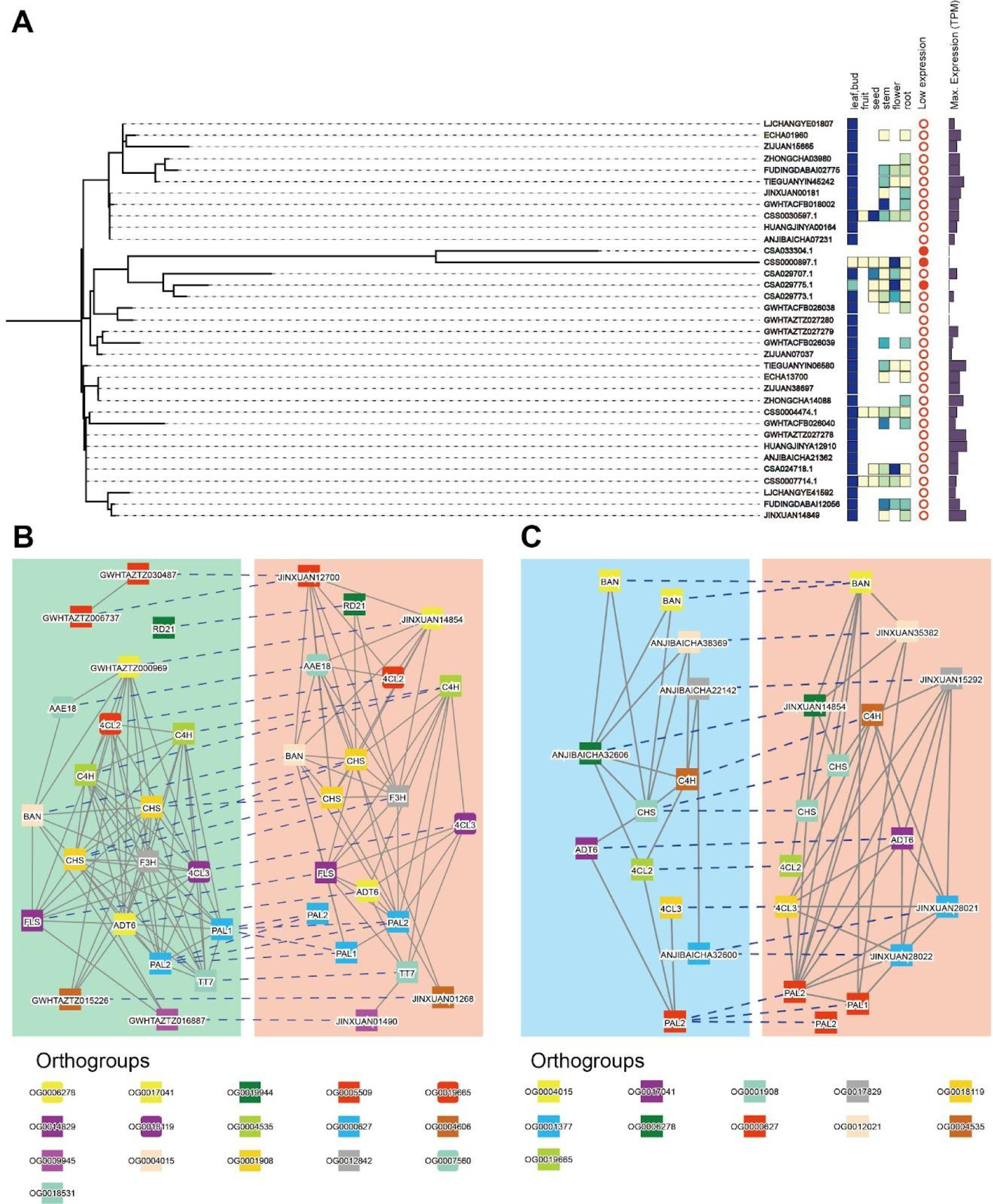
Comparative analysis of the CHS-related co-expression cluster among *Camellia sinensis* cultivars. (A) Phylogenetic tree of the OG0001908 orthogroup. The heatmap shows the expression level in different tissues, full red dots show genes with low-expression and the bar on the right indicates the maximum expression level (TPM). The color of a gene identifier indicates the cultivars. Note that OrthoFinder tree nodes do not contain bootstrap values, and should be interpreted with care. Missing data is indicated by an absent box. (B) Comparative analysis of the CHS-related co-expression cluster between “Huangdan” and “Jinxuan”. (C) Comparative analysis of the CHS-related co-expression cluster between “Anji Baicha” and “Jinxuan”.

To investigate the conservation of co-expression relationships within the chalcone synthase (CHS) gene family, we used TeaNekT comparative analysis. We compared “Cluster_126” of “Jinxuan” and “Cluster_216” of “Huangdan”, and observed that most of the flavonoid pathway-related gene families in “Cluster_126” are conserved between the two cultivars, such as PAL, C4H, 4CL, CHS, F3H and FLS gene families (https://teanekt.sbs.ntu.edu.sg/graph_comparison/cluster/2793/7475/1) (Figure 6B). Similarly, compared “Cluster_126” of “Jinxuan” and “Cluster_232” of “Anji Baicha”, and observed that most of the flavonoid pathway-related gene families in “Cluster_126” are conserved between the two cultivars, such as PAL, C4H, 4CL and CHS gene families (https://teanekt.sbs.ntu.edu.sg/graph_comparison/cluster/2106/7475/1) (Figure 6C). Intriguingly, we discovered three genes *JINXUAN35382*, *JINXUAN01268* and *JINXUAN01490* are involved in the co-expression network of “Cluster_126” in “Jinxuan” and exhibit conservation (Figure 6BC). As all three genes have a biological function labeled as “Not assigned”, this indicates that further research is needed to understand their specific biological functions (Figure 5C).

## Discussion

This study primarily conducted two aspects of work. Firstly, transcriptome assembly and comparative analysis were performed on 13 *Camellia sinensis* cultivars. Secondly, annotation, phylogenetic analysis, and co-expression network construction were carried out on the coding sequences of 13 cultivars, which were imported into the CoNekT framework to generate the TeaNekT database.

In previous studies, researchers tended to consider the *Camellia sinensis* species as a whole when conducting large-scale co-expression analysis of tea plant samples (Zhang et al., 2020). They generally used the genome of the *Camellia sinensis* cultivar “Shuchazao” as the reference genome to calculate gene expression levels for constructing the co-expression network of tea plants. In this study, the transcriptome of different *Camellia sinensis* cultivar was separately assembled to obtain their respective coding sequences. Each cultivar possesses unique coding sequences, allowing us to explore gene conservation through sequence similarities. Moreover, by analyzing these distinctive coding sequences, we can uncover cultivar-specific genes. For instance, during our comparative analysis of coding sequences from 13 cultivars, we observed that although the enriched biological functions of conserved genes are generally conserved across most cultivars, some variations still exist, such as the “transcription factor (ERF) activities” in “Yunkang 10” and the “FRS/FRF-type transcription factor” in “Zhongcha 108” (Figure S3A). Regarding the comparison of cultivar-specific genes among the coding sequences of the 13 cultivars, we discovered that “oxidative phosphorylation” in “Zijuan” and “phytohormone action.signalling peptides” in “Zhongcha 108” were specifically enriched (Figure S3B). Moreover, in the comparative analysis of transcription factor families among the coding sequences of the 13 cultivars, we observed that C2H2, MYB, bHLH, AP2/ERF-ERF, and NAC exhibited larger gene family sizes in almost all cultivars (Figure S4A). However, there were some exceptions, such as the relatively smaller C2H2 gene family in “Yunkang 10”.

Additionally, assembling the cultivar-specific transcriptomes provides us with an approach to study the phylogenetic relationships among tea cultivars. Such phylogenetic analysis can assist us in inferring the relationships between cultivars, enabling us to demonstrate the trends in differential formation during comparative analyses (Figure 4BC; Figure S3; Figure S4). For instance, we found that conserved genes were not enriched in the biological function “EC_4.2 carbon-oxygen lyase” of the two cultivars of var. *assamica*, while they were enriched in the biological function “EC_4.2 carbon-oxygen lyase” of the cultivars of var. *sinensis* (Figure S3A).

To give access to the newly assembled transcripts and gene expression data of various tea cultivars, we constructed the TeaNekT database. The database provides a comparative analysis of gene expression profiles, enabling researchers to identify factors influencing the expression levels of a group of genes (Figure 5A). The database provides advanced comparative expression profile and co-expression analyses and annotates the biological functions of the cluster, facilitating the identification of unique and conserved genes related to specific biological functions (Figure 6BC). By combining phylogenetic trees and tissue expression profiles, the tool allows researchers to understand the conservation of genes across tissues and identify tissue-specific genes (Figure 6A).

The usage of TeaNekT was exemplified by analyzing the co-expression cluster “Cluster_126” of “Jinxuan” cultivar. We observed that most members of the phenylalanine ammonia-lyase (PAL), cinnamate 4-hydroxylase (C4H), 4-coumarate-CoA ligase (4CL), and chalcone synthase (CHS) gene families show conserved co-expression across various cultivars. In addition, we observed several conserved, uncharacterized genes (Figure 6BC). Since conserved genes are likely to be functionally relevant (Hansen et al., 2014), we envision that further comparative analyses will uncover more genes important for the different characteristics of tea cultivars.

## Supporting information

Figure S1. Pseudoaligned reads percentage and sequencing reads distribution of the Camellia sinensis RNA-Seq samples.

Figure S2. Length distribution of coding sequences in the transcriptome of Camellia sinensis cultivars.

Figure S3. Functional enrichment heatmap of conserved genes and cultivar-specific genes using Mapman annotations as biological functions.

Figure S4. Copy number and median coefficient of variation (CV) of expression levels among transcriptional factor families in Camellia sinensis.

Table S1. Annotation information for RNA-Seq samples.

Table S2. Transcriptome assembled from 13 Camellia sinensis cultivars contain all coding sequences.

Table S3. Annotation information for 13 Camellia sinensis cultivars genes.

Table S4. Functional enrichment analysis of conserved genes and cultivar-specific genes using Mapman annotations as biological functions.

## Acknowledgments

X.Z. is sponsored by a China Scholarship Council fellowship.

## Author contributions

X.Z. led the main work of this study, including project conception, data annotation, data analysis, and paper writing. P.K.L. provided X.Z. with suggestions on transcriptome assembly. Z.A. uploaded the transcriptome data to CoNekT to create TeaNekT. M.M. and Y.W. co-supervised X.Z. in completing this project. M.M. and P.K.L. both participated in the revision of the paper. The authors thank all members of the Mutwil Lab for their suggestions and assistance with this manuscript.

## Supplementary tables

**Table S1. Annotation information for RNA-Seq samples, including tissue, sampling age, experimental treatment, processed reads number (NPR) and pseudoaligned rate (PPR).** (A-M) are “Anji Baicha”, “Echa 1”, “Fuding Dabaicha”, “Huangdan”, “Huangjinya”, “Jinxuan”, “Longjing 43”, “Longjing Changye”, “Shuchazao”, “Tieguanyin”, “Yunkang 10”, “Zhongcha 108”, “Zijuan”, respectively.

**Table S2. Transcriptome assembled from 13 *Camellia sinensis* cultivars contain all coding sequences.** (A-M) are “Anji Baicha”, “Echa 1”, “Fuding Dabaicha”, “Huangdan”, “Huangjinya”, “Jinxuan”, “Longjing 43”, “Longjing Changye”, “Shuchazao”, “Tieguanyin”, “Yunkang 10”, “Zhongcha 108”, “Zijuan”, respectively.

**Table S3. Annotation information for 13 *Camellia sinensis* cultivars genes, including ITAK annotation, KEGG annotation, GO annotation, Pfam annotation, and Mapman annotation, standard deviation (SD), coefficient of variation (CV) of expression levels and conservative or specific.** (A-M) are “Anji Baicha”, “Echa 1”, “Fuding Dabaicha”, “Huangdan”, “Huangjinya”, “Jinxuan”, “Longjing 43”, “Longjing Changye”, “Shuchazao”, “Tieguanyin”, “Yunkang 10”, “Zhongcha 108”, “Zijuan”, respectively.

**Table S4. Functional enrichment analysis of conserved genes and cultivar-specific genes using Mapman annotations as biological functions.** (A) FDR values for conserved genes. (B) Recalls for conserved genes. (C) FDR values for cultivar-specific genes. (D) Recalls for cultivar-specific genes.

## Supplementary figures

**Figure S1. Pseudoaligned reads percentage and sequencing reads distribution of the *Camellia sinensis* RNA-Seq samples.** The color of the points in the scatter plot represents the number of pseudoaligned reads.

**Figure S2. Length distribution of coding sequences in the transcriptome of *Camellia sinensis* cultivars.**

**Figure S3. Functional enrichment heatmap of conserved genes and cultivar-specific genes using Mapman annotations as biological functions.** The color of the cells represents recalls. The sorting of biological functions is based on the hierarchical clustering of recalls, while the sorting of cultivars is based on the phylogenetic tree. (A) Conserved genes. (B) Cultivar-specific genes.

**Figure S4. Copy number and median coefficient of variation (CV) of expression levels among transcriptional factor families in *Camellia sinensis*.** (A) Copy number. The color of the cells represents the copy number. The sorting of biological functional terms is based on the hierarchical clustering of the copy number, while the sorting of cultivars is based on the phylogenetic tree. (B) Median coefficient of variation (CV) of expression levels. The color of the cells represents the median coefficient of variation (CV) of expression levels. The sorting of transcription factor families is based on the hierarchical clustering of the median coefficient of variation (CV) of expression levels, while the sorting of cultivars is based on the phylogenetic tree.

