## Supplementary figures and images for "Comparative transcriptome database for *Camellia sinensis* reveals genes important for flavonoid synthesis in tea plants"

### Figure S1. Pseudoaligned reads percentage and sequencing reads distribution of the Camellia sinensis RNA-Seq samples.

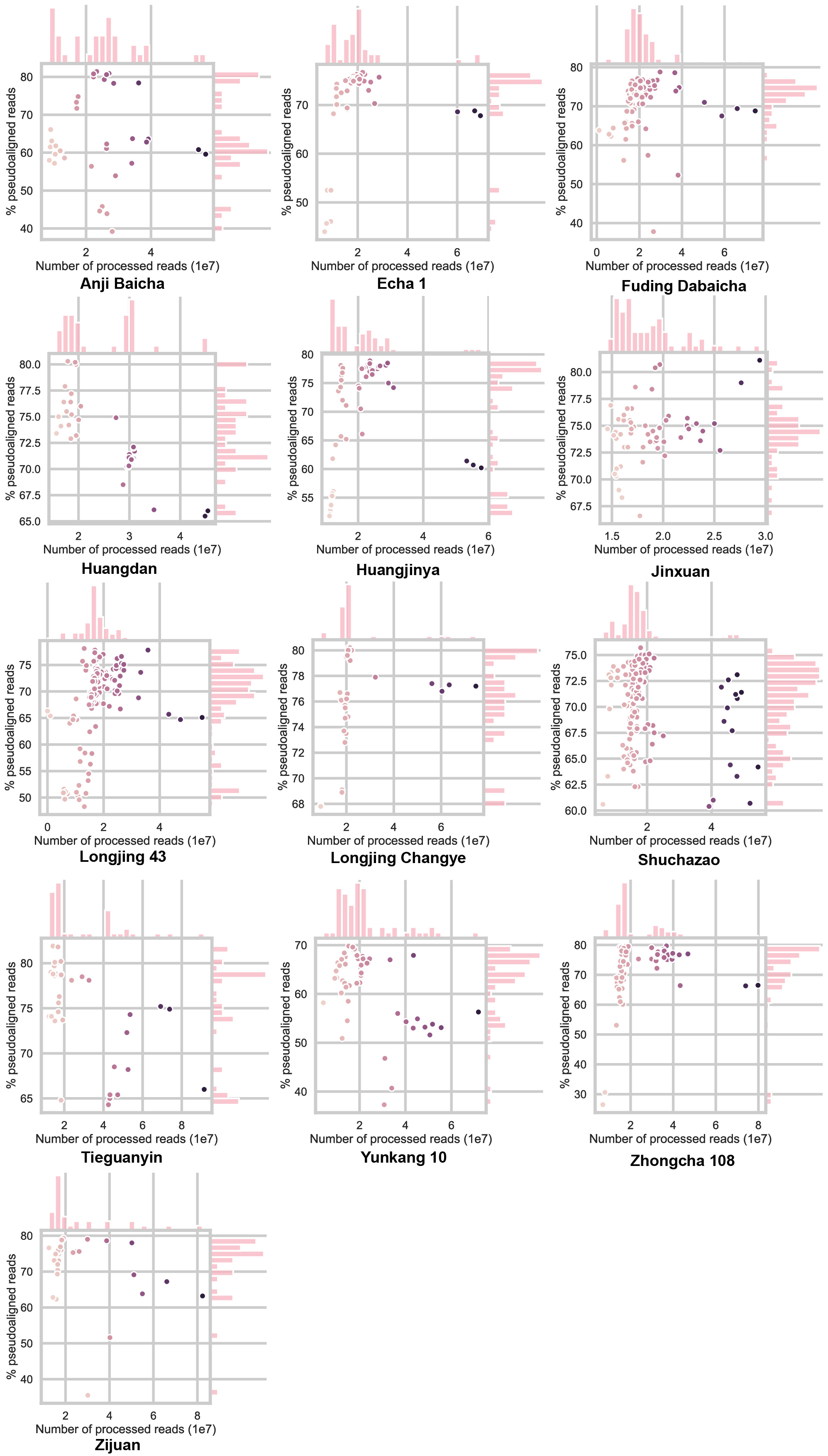

### Figure S2. Length distribution of coding sequences in the transcriptome of Camellia sinensis cultivars.

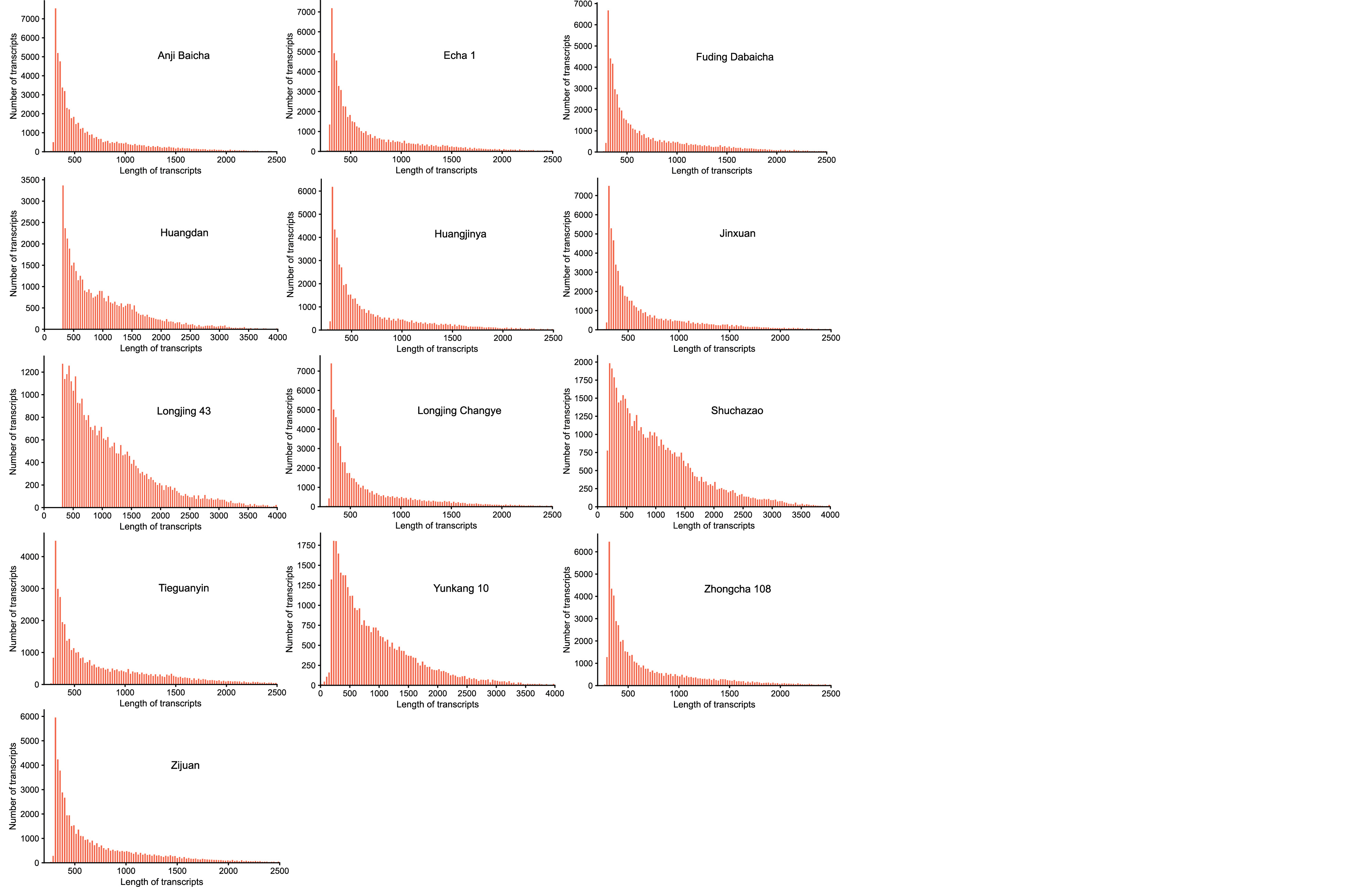

### Figure S3. Functional enrichment heatmap of conserved genes and cultivar-specific genes using Mapman annotations as biological functions.

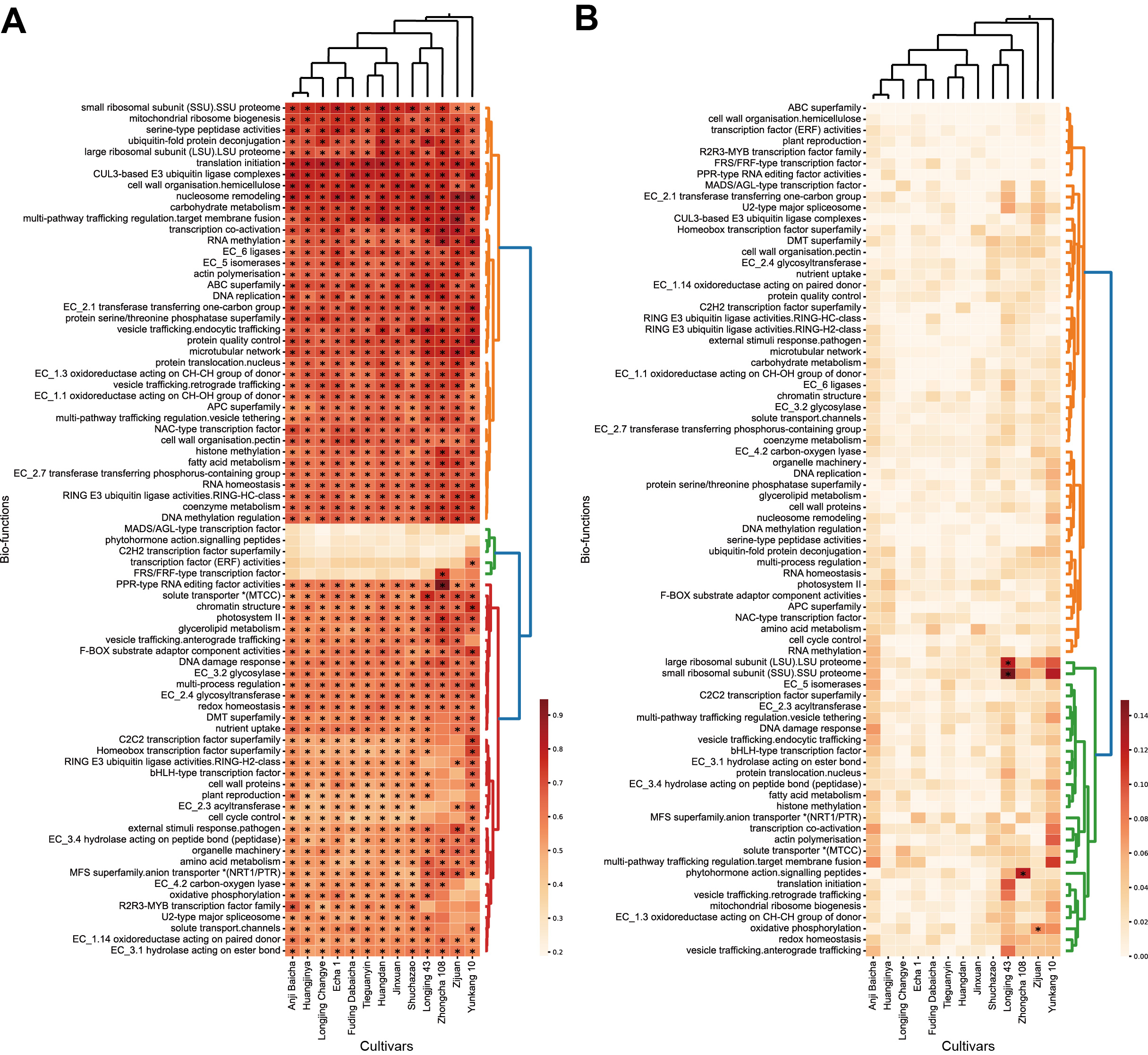

### Figure S4. Copy number and median coefficient of variation (CV) of expression levels among transcriptional factor families in Camellia sinensis.

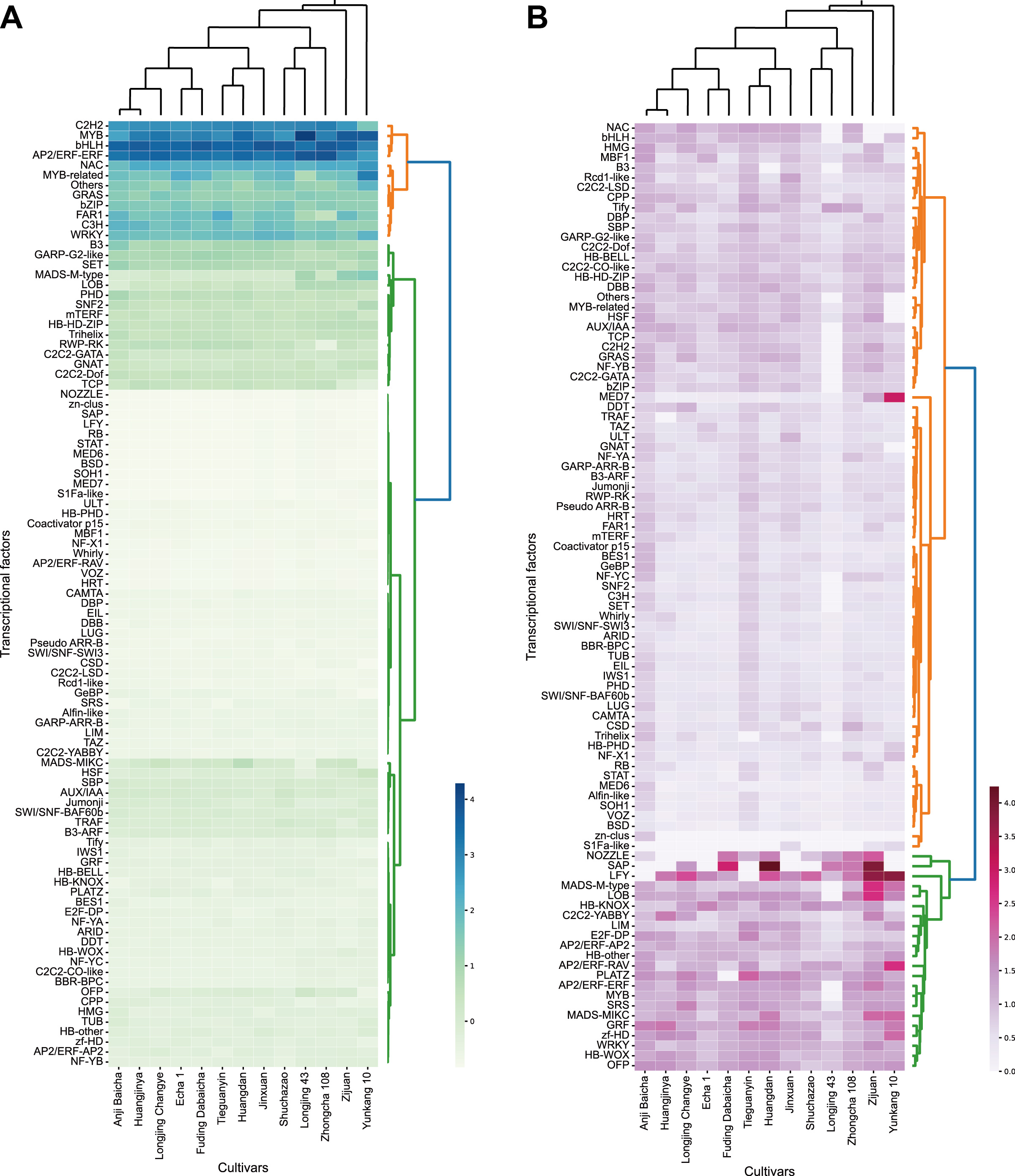
